# genBaRcode – a comprehensive R package for genetic barcode analysis

**DOI:** 10.1101/696229

**Authors:** Lars Thielecke, Kerstin Cornils, Ingmar Glauche

## Abstract

**Motivation:** Genetic barcodes have been established as an efficient method to trace clonal progeny of uniquely labeled cells by introducing artificial genetic sequences into the corresponding genomes. The assessment of those sequences, relies on next generation sequencing and the subsequent analysis aiming to identify sequences of interest and correctly quantifying their abundance.

**Results:** We developed the *genBaRcode* package as a toolbox combining the flexibility of digesting next generation sequencing reads with or without a sophisticated barcode structure, with a variety of error correction approaches and the availability of several types of visualization routines. Furthermore, a graphical user interface was incorporated to allow also less experienced R users package-based analyses. Finally, the provided tool is intended to bridge the gap between generating and analyzing barcode data and thereby supporting the establishment of standardized and reproducible analysis strategies.

**Availability:** The *genBaRcode* package is available at CRAN (https://cran.r-project.org/pack-age=genBaRcode).

## 1 Introduction

The ability to mark single cells and to follow their progeny is a key requirement for many biological experiments. During the last decade, genetic barcodes were introduced as a new tool to genetically label single cells and all their future descendants [1-4]. It quickly became the method of choice for the majority of cellular labeling studies. Consequently, research groups developed their own barcode constructs, including the selection of different viral vectors and amplification protocols [2, 3, 5-9]. Given the lack of standardization and available software solutions, specifically tailored, in-house solutions were used in order to analyze the resulting data. Unfortunately, those different analysis strategies harbor the risk of generating results that, at the very least, are difficult to compare. This general short-coming of standardized reporting and analysis pipelines was recently outlined as a major limitation to further expand the application of barcoding and clonal tracking experiments [10].

Encountering similar problems while developing our own barcode constructs [11, 12] and comparing our solutions to already published results, we decided to develop a comprehensive R-package combining basic functionalities for data extraction and analysis with a collection of visualizations wrapped in a set of simple albeit adjustable R-functions. By adding an ancillary *shiny-app* we also provide a graphical user interface (GUI) to make the package accessible not only to experienced R-programmers but also to investigators lacking the necessary programming skills.

We are convinced that such a “toolbox”-like software package will contribute to standardization efforts in the field of barcode analysis [10]. Therefore, we deliberately chose R as a platform, as it is one of the most wide-spread tools for data analyses utilized by both bioinformaticians and biologists. Furthermore, R is a freely available software that has already a large community constantly developing and publishing a large variety of additional functionality (CRAN [13], Bioconductor [14]). In the following we present the “*genBaRcode”*-package, currently at version 1.2.2, and describing an exemplary work flow spreading from data extraction to data visualization.

## 2 Implementation

The package is available at CRAN and can be installed like every other CRAN R-package:

~~~
> *install.packages(“genBaRcode”)*
~~~

Unfortunately, while installing CRAN packages it is not always possible to automatically install also the necessary Bioconductor packages. Therefore, if the installation process ends with an error message similar to “*ERROR: dependencies ‘Biostrings’, ‘ShortRead’, ‘ggnetwork’, ‘S4Vectors’, ‘ggtree’ are not available for package ‘genBaRcode’*”, the user has to install those packages manually:

~~~
> *if (!requireNamespace(“BiocManager”, quietly = TRUE)) {install.packages(“BiocManager”)*
}
> *BiocManager::install(c(“Biostrings”, “ShortRead”, “S4Vectors”, “ggtree”))*
~~~

After this installation process, the package can simply be loaded and used:

~~~
> *library(“genBaRcode”)*
~~~

### 2.1 Data extraction and analysis

The first part of the package is dedicated to the initial data processing. It provides the function *processingRawData()*, which conveniently combines all necessary steps for the translation of raw next generation sequencing (NGS) data files into a sorted table of barcode (BC) sequences and its corresponding read counts.

~~~
> *bb <-getBackboneSelection(1)*
> *s_dir <-system.file(“extdata”, package = “genBaRcode”)*
> *res_dir <-“/my/results/directory/”*

> *BC_dat <-processingRawData(file_name = “test_data.fastq.gz”,
source_dir = s_dir, results_dir = res_dir, mismatch = 1, label = “test”,
bc_backbone = bb, min_score = 30, min_reads = 2, save_it = FALSE,
seqLogo = FALSE, cpus = 4, wobble_extraction = TRUE,
dist_measure = “hamming”)*
~~~

Technically, the user has to supply one or multiple NGS data files (*file_name*) and the path to the corresponding directory (*source_dir*). Within the current version FASTA as well as FASTQ file formats will be accepted. Furthermore, the user can provide none, one or multiple BC design pattern(s), so-called BC-backbones (*bc_backbone*), which, if supplied, are then used as identifier for the actual BCs within the individual NGS reads. A BC-backbone needs to be encoded as a character string containing ‘N’s at the actual BC positions (also called wobble positions) and nucleotides (A, T, G, C) at the accompanying fixed positions. Those fixed positions are essential in order to identify individual BC sequences by matching them against the actual read. An exemplary backbone could look like this: *“ATCNNTAGNNTCCNNAAGNN”*. The matching procedure also allows to specify a number of acceptable sequence mismatches (*mismatch*). Moreover, if the parameter *wobble_extraction* was set to *TRUE*, the identified sequences will be stripped from their fixed backbone positions and only the actual BC sequences (wobble positions) will be sustained. Otherwise only the flanking regions of the read will be stripped and the identified BC including the wobble positions will be returned. In case no BC backbone was chosen, all NGS reads will be treated as BCs and clustered based on the declared number of accepted dissimilarities (*mismatch*) and the elected distance measure (*dist_measure*).

Furthermore, there are additional possibilities to customize the analysis procedure. In case a FASTQ data file is available, the *min_score* parameter restricts the analysis to reads with a certain minimum sequence quality (calculated as the Phred-score average over the entire read length). Optionally, the quantified BC sequences can automatically be filtered for a minimum number of reads (*min_reads*) to exclude contaminations or untrustworthy reads. A sequence logo of the entire NGS data file can be created (*seqLogo*) and a csv-file of all detected BCs including their corresponding read counts can be saved (*save_it*). Both will be stored within the specified results directory (*results_dir)* for all the subsequent analyses.

In case of only one specified BC-backbone, only one csv-file will be created while the function returns a specifically designed S4 data object (called *BCdat* data type) containing not only the detected BCs but also important parameters such as the used backbone pattern, a unique label used as name tag e.g. for all saved csv-files and the path to the results directory. If multiple backbones and/or NGS files are provided, also multiple csv-files will be created while the returned object will be a list containing several similar S4 data objects in the same order as the provided NGS files and BC-backbones.

The user can also decide, if available, how many CPUs (*cpus*) should be utilized. The algorithm will then automatically start the analyses in a parallelized fashion, distributing not only the BC extraction but also the parallel analysis of multiple NGS data files onto multiple cores. Providing several BC-backbones at once will shorten the computational time compared to individual function calls with only one given BC-backbone each. Admittedly, the algorithm sequentially searches for a given BC-backbone but since the read length typically only allows for just one BC construct per NGS read, it will automatically dismiss reads with an already identified BC and therefore significantly reduce the search space for the next BC-backbone(s). Furthermore, since the parallelization is based on the *future* R-package, it is also possible to choose a future-specific strategy in order to utilize the capacity of different resources in the best way possible.

The entire package is suitable for any kind of BC designs as long as they adhere to the aforementioned backbone structure. The BC extraction function is based on the “*Biostrings”*-package, meaning that the BC design pattern will not only allow for the wildcard letter ‘N’ but also for all letters which are part of the nucleic acid notation of the International Union of Pure and Applied Chemistry (IUPAC notation), making it highly customizable and capable of searching for almost every kind of BC design. Additionally, utilizing the *getBackboneSelection()* function, the user can select from eight predefined and already published BC designs [11, 12].

After the BC identification, the newly created *BCdat* object(s) can then be subjected to the following function to initiate the error-correction (EC) of the detected BCs.

~~~
> *BC_dat_EC <-errorCorrection(BC_dat, maxDist = 8, save_it = FALSE, m = “hamming”, type = “standard”, EC_analysis = FALSE)*
~~~

This default EC-procedure (*type = “standard”*) is an implementation of the method described in Thielecke et al. [11] that merges highly similar BC sequences in order to reduce erroneous/diverging BC sequences, which regularly occur as a result of polymerase chain reactions (PCR) during sample preparation and NGS. Sequence similarity calculations are based on a given distance measure *m.* For equally sized BCs we would recommend using the *Hamming* distance, but the package is also able to calculate a broad variety of additional measures, e.g. the Levenstein distance (*m = “lv”*), the longest common substring distance (*m = “lcs”*) or the Jaro-Winker distance (*m = “jw”*). The following command will show the user all available distance measures, provided by the *stringdist()* function of the *stringdist* package.

~~~
> ?’stringdist-metrics’
~~~

All of those sequence similarity calculations are further accompanied by an adjustable similarity threshold *maxDist*. BCs with a distance < *maxDist* will be clustered together, starting with the least frequent BC. In case there is more than one BC with the same distance, per default, only the two BCs with the lowest read count will be clustered together (the method is inspired by the idea that during PCR successive point mutations occur and, in a perfect world, one would find a chain of BCs successively increasing the number of diverging nucleotides while decreasing the number of read counts).

Although the *“standard”* method is recommended by us, the parameter *start_small*, if set to FALSE, offers the possibility to invert the procedure and therefore to always cluster the BC of interest starting with the most frequent one.

There is also the EC-type *“connectivity based”* available, which follows the *“standard”* procedure but instead of ordering the available BCs by read counts and starting the clustering with the least frequent one, it now will be ordered by the amount of highly similar analogues (depending on the *maxDist* value) and consequently the clustering will start with the BC possessing the lowest number of highly similar counterparts.

The *“graph based”* EC-type is based on the idea of graph-theoretic (connected) components. Firstly, for each and every BC the distances to all the other BCs will be calculated and all distances > *maxDist* will then be set to zero. Secondly, the resulting matrix will serve as an adjacency matrix for which existing components can be identified. Finally, all of the “member-BCs” of those components will be clustered together, with the most abundant BC as the respective “cluster-label”.

The EC-type called *“clustering”* just starts with the most frequent BC, identifies all highly similar counterparts (again based on the method *m* and the threshold *maxDist*), adds up all of the corresponding reads and lastly all of those added up BC sequences will be dismissed. Then, the procedure continues with the second most abundant BC until all BCs are processed. Since the actual EC-algorithms are hardly parallelizable, parallel computations are only feasible if a list of *BCdat*-objects is supplied. The *error-Correction()* routine will automatically initiate a parallel execution depending on the number of data objects and the *cpus* parameter. The resulting data object will again be one or a list of *BCdat* objects. Optionally, the final list of corrected BC sequences can also be saved as a common csvfile (*save_it*) in the specified directory (*results_dir*).

### 2.2 Data visualization

The second part of the package covers the data visualization. The implemented routines also include illustrations to check the quality of the supplied NGS data. As such, it is possible to visualize the frequency of mean quality scores over all reads (Fig. 1a) and an overview plot which displays the quality score distribution per sequencing cycle (Fig. 1b). Additionally, the function *plotSeqLogo()* creates a sequence logo either from all NGS reads (see *processingRawData()*), the entire barcode construct or only the wobble positions.

**Figure 1.**
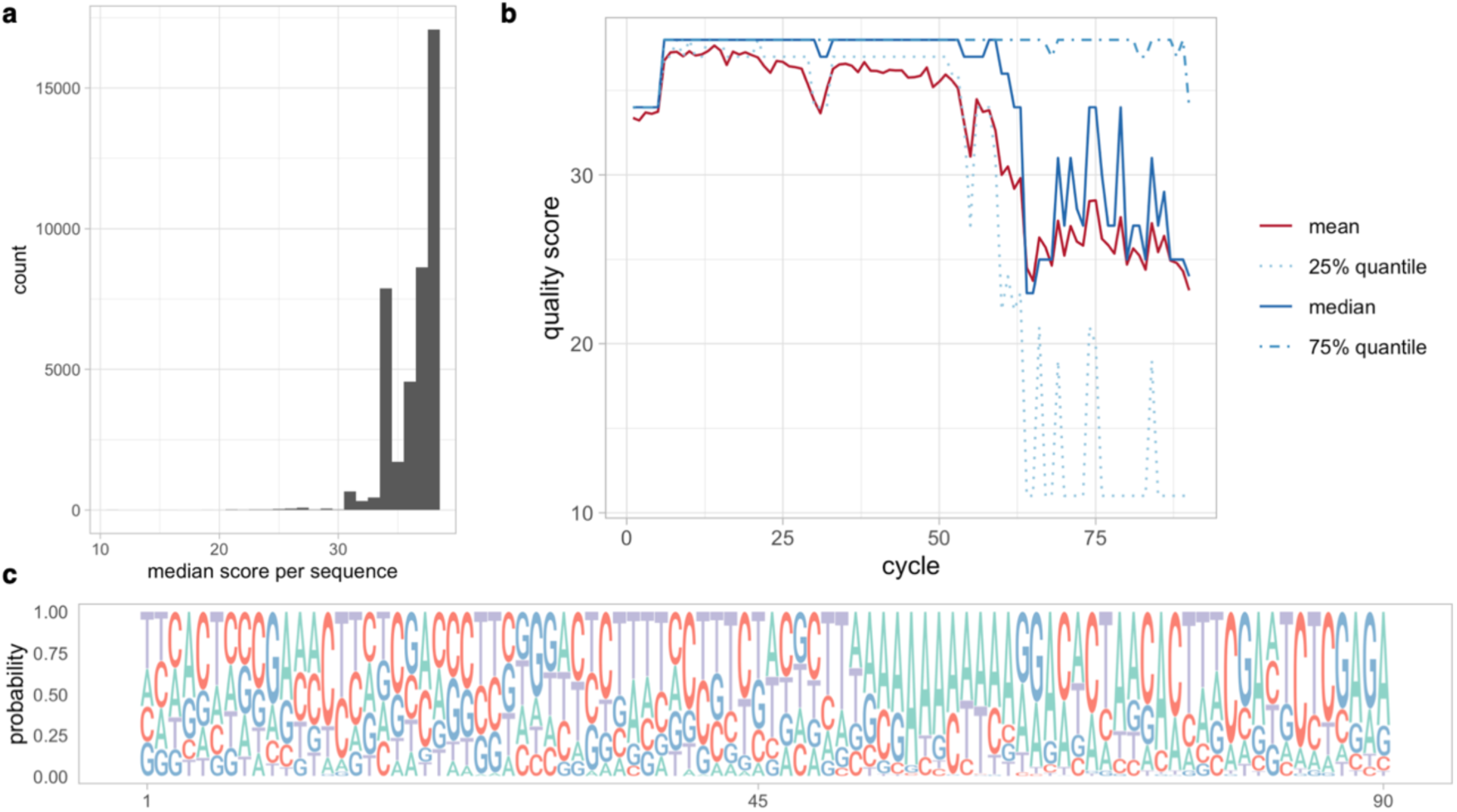
The quality of NGS data files can be visualized by plotting **(a)** the frequency of the mean quality scores over all reads, **(b)** the mean and median quality scores per sequencing cycle and/or **(c)** the sequence logo of the whole NGS file over the entire read length.

Such plots offer the possibility to visually check for prominent mismatch positions within the backbone or general irregularities regarding the nucleotide distribution (Fig. 1c).

To examine the actual BC data, the plot function *generateKirchenplot()* depicts the sorted read counts of all detected BCs (before or after error-correction) as a simple barplot (Fig. 2a).

**Figure 2.**
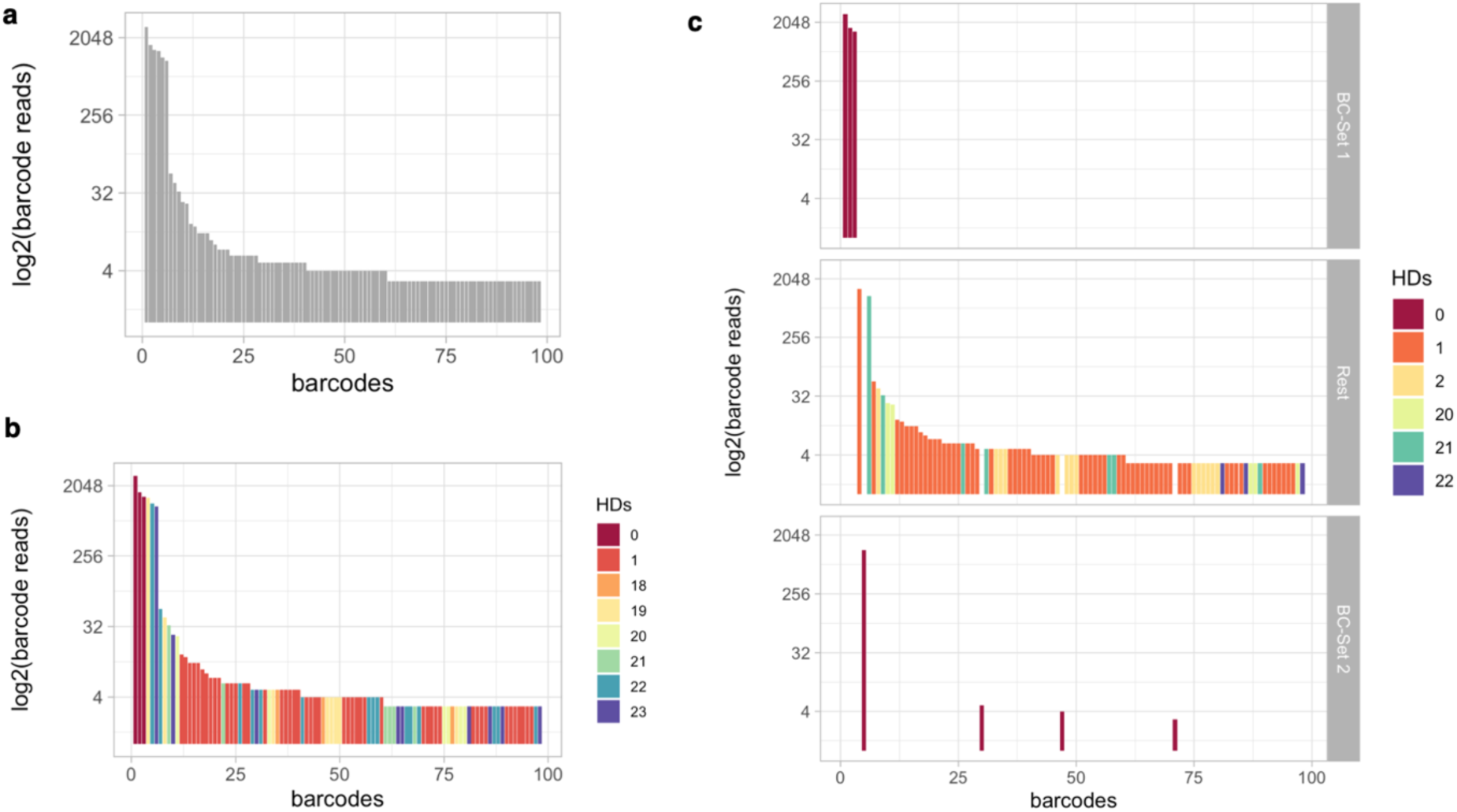
A simple barplot of sorted read counts of all detected BCs **(a)** will be generated by the function *generateKirchenplot()*. If the user also provides a list of known reference BCs **(b)** there will be a color-coding of each detected BC according to its sequence similarity value of the most similar reference BC. (**c**) If, at the same time, two distinct subsets of known BCs are provided there will be three panels containing the respective BC subsets.

~~~
> *generateKirchenplot (BC_dat, ori_BCs = NULL, ori_BCs2 = NULL, loga = TRUE, col_type = NULL, m = “hamming”)*
~~~

If the user is interested in a subset of known BCs, a corresponding list of those reference BCs can be provided (*ori_BCs*), thereby enabling a color-coding of each measured BC according to its minimal sequence similarities to the entire subset of reference BCs (Fig. 2b). It is also possible to provide two distinct subsets of known BCs (*ori_BCs, ori_BCs2*), e.g. BCs of the designed BC library and known contaminations (Fig. 2c). To make a visual discrimination of those two subsets easier, the resulting plot consists of three panels containing all BCs of the respective subset. By default, the function calculates the *Hamming* distances (HD, number of nucleotide differences) between the reference set and all other BCs and uses the minimum HD as color-coding basis. The information about sequence similarities (before EC) can be used to either approximate the extend of generated sequence errors (assuming that highly similar sequences are caused by one or few single nucleotide exchanges) or to visually check the uniqueness/discriminability of the used BCs in terms of total sequence variations (after EC).

The sequence similarities can also be conveniently visualized in a network-like structure. Figure 3 shows one instance of a possible variety of different network plots utilizing different third-party R-packages (*ggnetwork: ggplotDistanceGraph(), igraph: plotDistanceIgraph(), visNetwork: plotDistanceVisNetwork()*) available. This kind of visualization represents every BC sequence as a node and every edge indicates a minimal similarity value between adjacent nodes or BC sequences (by default a minimal Hamming distance of *m* = 1 will be visualized). It is also possible to connect each node with its most similar neighbor (*complete*), assuring at least one edge per node and therefore allowing for Hamming distances > 1. While these plots are 2D representations, utilization of the “*rgl”*-package allows for the creation of an interactive 3D graph (encoded by the *threeD* parameter of the *plotDistanceIgraph*() function). As further options, the user can also create a *gdf*-file (*createGDF()*), which was defined and is used by GUESS (Graph Exploration System) but can also be imported into the open source graphic software *Gephi* [15]. Both programs are specialized in visualizing networks and allow for an even broader variety of design options.

**Figure 3.**
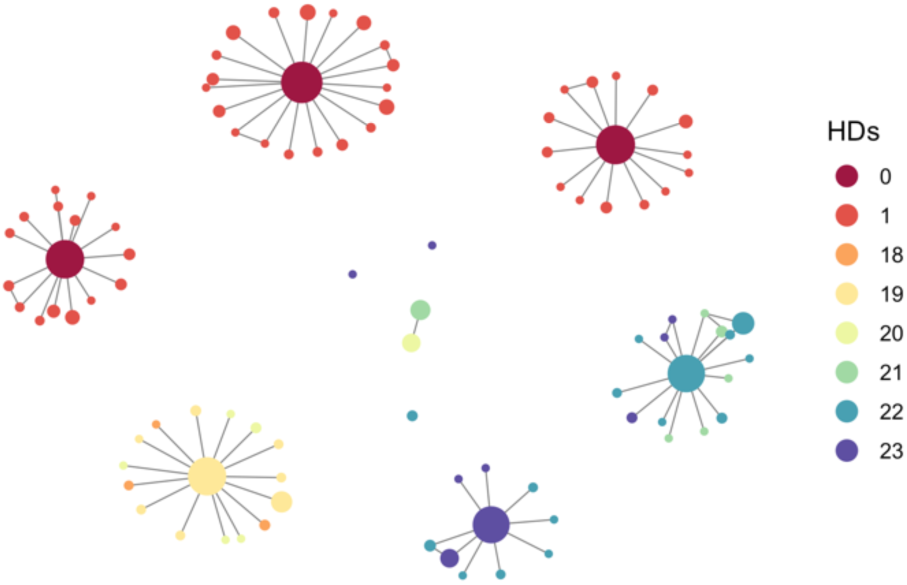
The sequence similarities can also be visualized in a network- or tree-like structure. Within a network-based visualization, every BC sequence will be represented as a node and every edge indicates a minimal similarity value between adjacent nodes (BC sequences). By default, a minimal Hamming distance of 1 will be visualized but it is also possible to always connect each node with its most similar neighbor, therefore assuring that each node will be connected to at least one other node (allowing for a possible hamming distance of >1).

Besides a network-based representation it is also possible to visualize sequence similarities as a tree-like structure utilizing the *plotClusterGgTree()* function. In those plots, each forked branch and its respective distance to the previous branch represents the estimated sequence relationship.

Additionally, it is also possible to visualize the EC-procedure in order to develop an understanding of the amount and specificity of the corrected errors or/and to decide which EC-approach best suits the specific data set. In order to generate the needed data, the *errorCorrection()* function provides the parameter *EC_analysis* which, if set to TRUE, will document the entire EC-process (since a lot of additional data will be calculated and stored, it will slow down the execution time noticeable, therefore it is recommended to activate it only when needed). The returned *BCdat* object will then be composed not only of the usual BC list but will also include EC-specific cluster information (i.e. cluster sizes, added read counts, cluster specific BC lists) necessary for the aforementioned EC-plots.

Depending on the EC-method the composition of single clusters is not always foreseeable therefore the function *error_correction_clustered_ HDs()* will create a scatter plot displaying the maximal distances within each of the barcode sequence clusters. Furthermore, the two functions *error_correction_circlePlot()* and *error_correction_treePlot()* will both create a specialized plot visualizing the size of the emerged barcode clusters including the abundance of the involved BCs. Both functions visualize the same kind of data but one in a circular and the other in a tree-like representation style.

The “*genBaRcode”*-package also offers the possibility to analyze and visualize time series data. As already mentioned, the *processingRawData()* function also accepts a list of NGS files which if *cpus* > 1 will be analyzed in parallel and will result in a list of *BCdat* objects. In order to analyze repetitive measurements, *generateTimeSeriesData()* takes such a list of *BCdat* objects and calculates a matrix containing all unique BC sequences over all single *BCdat* objects as rows and their respective number of read counts over all time points (data objects) as columns. The *plotTimeSeries()* function utilizes such a matrix to depict the time dependent development of each and every BC detected at least once. Furthermore, it is also possible to provide a vector containing the time information of each measurement which will then be factored in the final plot to achieve the appropriate time scale.

Finally, the package also offers the option to visualize multiple measurements as Venn diagrams. The function *plotVennDiagramm()* also accepts a list of *BCdat* objects from which the corresponding Venn diagram will be generated. Almost all of the mentioned plotting functions return a *ggplot2* object which can be plotted directly or customized further.

### 2.3 Graphical User Interface

In addition to the described functionality of the R-package, we developed an easy-to-use interface mainly for investigators with little or no programming experience. Therefore, the third and final piece of the package consists of a *shiny-app* making the majority of the aforementioned functions available within a graphical user interface (GUI). The GUI enables the unexperienced user to perform the majority of the available analyses essentially without typing any line of R-code. The app not only allows for the BC extraction from raw NGS data files, the error-correction and the storage of those BC sequences within external csv-files but also for a visual inspection of the corresponding results directly within the app and to further manipulate and save the generated figures. Furthermore, for the more skilled user the app doubles as a convenient possibility to familiarize oneself with the package’s functionality and its intended usage. It can easily be started utilizing the *genBaRcode_app()* function, immediately prompting the user to specify a path to the NGS files of interest. If no specific path is provided, the included template will be made available for example analyses.

Within the app, the user can specify one or multiple NGS files for a quantitative analyze and adjust further essential parameters (Fig. 4a). It is also possible to just read one or multiple existing csv-file(s) containing already extracted and counted BC sequences in order to generate new plots or to reanalyze certain data sets. If multiple files have been chosen, the app automatically initiates a time series analysis. BC extraction and error-correction will be performed according to the user specifications. After finishing the necessary calculations, a list of selectable plots and supplementing tabs containing meta-information as well as lists of the detected barcodes in combination with their respective read-counts before and after error-correction will become available (Fig. 4b).

**Figure 4.**
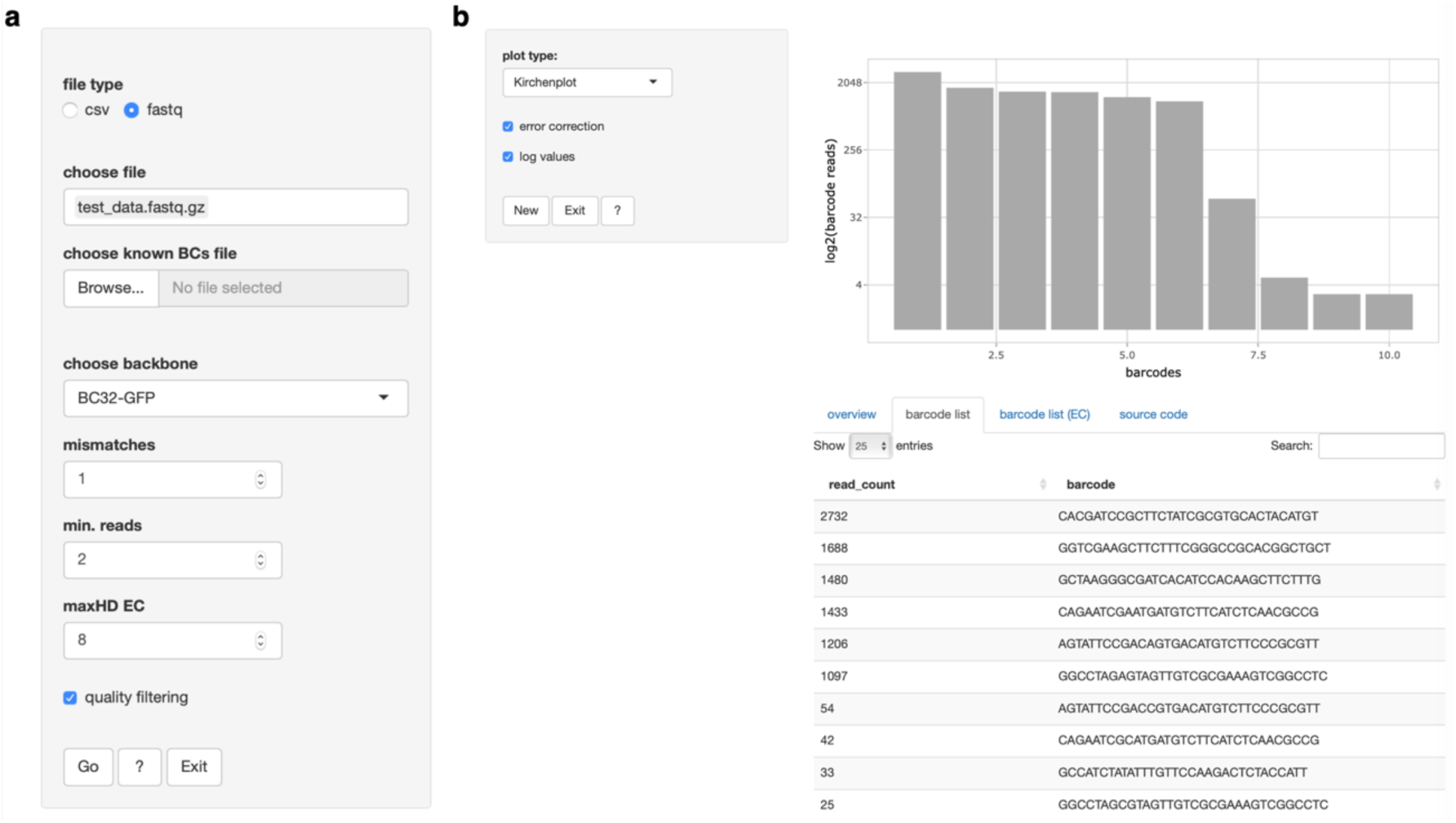
The included app allows for **(a)** a selection of one or multiple NGS or csv files for subsequent analyses and also for the specification of further essential analysis parameters. After finishing the necessary calculations, **(b)** a list of selectable plots and supplementing tabs containing meta-information will be available.

Selecting a particular plot style prompts the app to display the corresponding figure utilizing the *shiny* specific output und render function of the “*plotly”*-package, allowing not only an inspection but also an interaction with the produced figures via clicking, hovering and/or dragging. The user can freely zoom in and out, scroll through the image and extract additional information by hovering over certain elements of interest, thereby providing the functionality to intuitively explore the results and customize the respective visualizations.

In order to augment the support for less experienced users, an additional tab provides all the corresponding R-code, necessary to reproduce the presented results and figures within the command-line interpreter and without the assistance of the shown GUI. Furthermore, there is a detailed help-page available, designed to guide the user step-by-step through the app and also to make the basic R help accessible in order to further familiarize the user with the underlying R-functions.

Additionally, for a more in-depth tutorial and a step-by-step description the package is also equipped with so-called vignettes. Two separate documents demonstrating and describing the package functions and the graphical user-interface. They can either be found directly on CRAN [16] or can conveniently be generated directly within R after installing the package. The following command will reveal both available vignettes.

~~~
> browseVignettes(package = “genBaRcode”)
~~~

Furthermore, the source code of the entire package is available to the user on CRAN via GitHub [17] and all implemented unit tests can also be inspected directly within GitHub [18].

## 3 Discussion

The establishment of genetic barcoding and the availability of NGS boosted clonal studies in many areas of biology. This ultimately leads to an increasing demand for the analysis of the resulting BC data and calls for standardization efforts including BC calling, quantification, error-correction and visualization. Our R-package comprises currently available analysis routines and combines them with a user-friendly GUI. It is compatible with a variety of different BC constructs, published or newly developed, and it provides the necessary means to quantitatively analysis and visualize the resulting data obtained by NGS.

Furthermore, the GUI makes the analysis software accessible for investigators with only basic or no R-knowledge. Thereby our package bridges the gap between data generation and analysis, and supports interactions between investigators and data scientists, not only with respect to experiment-specific analysis pipelines, but also concerning the establishment of a consensual, standardized data processing and analysis strategy.

## Acknowledgements

The authors would like to thank Andreas Dahl, Sebastian Gerdes and Christoph Baldow for fruitful discussions concerning methods development and Ingo Roeder for providing the necessary infrastructure to perform this work. The authors also thank Sebastian Wagner for testing the software and suggesting improvements.

## References

1. Bystrykh LV, de Haan G, Verovskaya E: Barcoded vector libraries and retroviral or lentiviral barcoding of hematopoietic stem cells. Methods Mol Biol 2014, 1185:345–360.

2. Nguyen LV, Makarem M, Carles A, Moksa M, Kannan N, Pandoh P, Eirew P, Osako T, Kardel M, Cheung AM et al: Clonal analysis via barcoding reveals diverse growth and differentiation of transplanted mouse and human mammary stem cells. Cell Stem Cell 2014, 14(2):253–263.

3. Gerrits A. DB, Kalmykowa O. J., Klauke K., Verovskaya E., Broekhuis M. J. C., de Haan G., and Bystrykh L. V.: Cellular barcoding tool for clonal analysis in the hematopoietic system. Blood 2010, 115(13):2610–2618.

4. Perie L, Hodgkin PD, Naik SH, Schumacher TN, de Boer RJ, Duffy KR: Determining lineage pathways from cellular barcoding experiments. Cell Rep 2014, 6(4):617–624.

5. Peikon ID, Gizatullina DI, Zador AM: In vivo generation of DNA sequence diversity for cellular barcoding. Nucleic Acids Res 2014, 42(16):e127.

6. Lu R, Neff NF, Quake SR, Weissman IL: Tracking single hematopoietic stem cells in vivo using high-throughput sequencing in conjunction with viral genetic barcoding. Nat Biotechnol 2011, 29(10):928–933.

7. Nguyen LV, Pellacani D, Lefort S, Kannan N, Osako T, Makarem M, Cox CL, Kennedy W, Beer P, Carles A et al: Barcoding reveals complex clonal dynamics of de novo transformed human mammary cells. Nature 2015, 528(7581):267–271.

8. Cheung AM, Nguyen LV, Carles A, Beer P, Miller PH, Knapp DJ, Dhillon K, Hirst M, Eaves CJ: Analysis of the clonal growth and differentiation dynamics of primitive barcoded human cord blood cells in NSG mice. Blood 2013, 122(18):3129–3137.

9. Wu C, Li B, Lu R, Koelle SJ, Yang Y, Jares A, Krouse AE, Metzger M, Liang F, Lore K et al: Clonal tracking of rhesus macaque hematopoiesis highlights a distinct lineage origin for natural killer cells. Cell Stem Cell 2014, 14(4):486–499.

10. Lyne AM, Kent DG, Laurenti E, Cornils K, Glauche I, Perie L: A track of the clones: new developments in cellular barcoding. Exp Hematol 2018, 68:15–20.

11. Thielecke L, Aranyossy T, Dahl A, Tiwari R, Roeder I, Geiger H, Fehse B, Glauche I, Cornils K: Limitations and challenges of genetic barcode quantification. Sci Rep 2017, 7:43249.

12. Cornils K, Thielecke L, Huser S, Forgber M, Thomaschewski M, Kleist N, Hussein K, Riecken K, Volz T, Gerdes S et al: Multiplexing clonality: combining RGB marking and genetic barcoding. Nucleic Acids Res 2014, 42(7):e56.

13. The Comprehensive R Archive Network. https://cran.r-project.org. Accessed 08 Nov 2017

14. Bioconductor - Open Source Software for Bioinformatics. http://bioconductor.org. Accessed 08 Nov 2017

15. Bastian M, Heymann S, Jacomy M: Gephi: An Open Source Software for Exploring and Manipulating Networks; 2009.

16. genBaRcode - Vignettes, https://cran.rstudio.com/web/packages/genBaRcode/vignettes/, Accessed 25. Oct 2019

17. genBaRcode - Source Code, https://github.com/cran/genBaRcode/tree/master/R, Accessed 25. Oct 2019.

18. genBaRcode - Unit Tests, https://github.com/cran/genBaRcode/tree/master/tests/testthat, Accessed 25. Oct 2019.

